# Novel Abundant Oceanic Viruses of Uncultured Marine Group II Euryarchaeota Identified by Genome-Centric Metagenomics

**DOI:** 10.1101/101196

**Authors:** Alon Philosof, Natalya Yutin, José Flores-Uribe, Itai Sharon, Eugene V. Koonin, Oded Béjà

## Abstract

Marine Group II Euryarchaeota (MGII) are among the most abundant microbes in the oceanic surface waters. So far, however, representatives of MGII have not been cultivated, and no viruses infecting these organisms have been described. Here we present complete genomes for 3 distinct groups of viruses assembled from metagenomic sequence datasets highly enriched for MGII. These novel viruses, which we denote Magroviruses, possess double-stranded DNA genomes of 65 to 100 kilobase in size that encode a structural module characteristic of head-tailed viruses and, unusually for archaeal and bacterial viruses, a nearly complete replication apparatus of apparent archaeal origin. The newly identified Magroviruses are widespread and abundant, and therefore are likely to be major ecological agents.

---

Euryarchaeal Marine group II (MGII) members are highly abundant in the photic zone of oligotrophic oceans (*1-3*). This group has been reported to show seasonal variation (*4*) and to comprise up to 90% of the total archaea and one third of total microbial cells during spring blooms in the Atlantic Ocean (*5*). Metagenomic studies have demonstrated higher abundance of MGII in surface waters, mixed layer and halocline in the Arctic Ocean (*6, 7*), the Gulf of Mexico (*8*) and the Pacific Ocean (*9*). The MGII has been divided into two distinct sub-lineages, II.a and II.b (*10*). Metatranscriptomic analyses shows that the MGII archaea are among the most transcriptionally active microbial groups in the coastal Pacific Ocean, with transcription levels and patterns similar to those of *Pelagibacter ubique* and SAR86 (*11, 12*).

Despite their high abundance and transcription activity, to date, not a single representative of MGII has been cultured. Nevertheless, Iverson et al. (2012) (*13*) were able to assemble a complete MG II.a genome from metagenomic sequences. The analysis of this genome suggests that group MG II.a are photoheterotrophic (using proteorhodopsins) and motile (*14*). A representative genome from MG II.b (*Thalassoarcahea*) has been recently assembled from fosmid libraries derived from Deep Chlorophyll Maximum samples in the Mediterranean (*15*). Metagenomic fragment recruitment analysis indicates that MG II.a are more abundant in costal and estuarine habitats whereas MG II.b are more abundant in oligotrophic ocean habitats (*15*).

To gain further insight into the diel activity of MGII archaea in the Red Sea, we examined metagenomic bins containing MGII contigs. These bins were constructed from contigs retrieved from a cross assembly of microbial, viral and transcriptomic samples collected at four time points during a single day in the Gulf of Aqaba in the Red-Sea (ENA project PRJEB19060). Close manual inspection of these sequences showed that one bin (169) contained, along with contigs of MGII, a Metagenome-Assembled Genome (MAG) (156409) carrying hallmark viral genes including predicted Major Capsid Protein (MCP), Portal protein, and large subunit of the Terminase, as well as DNA Polymerase of the B family (DNAP). This contig was found to contain overlapping terminal regions suggesting that it represents a complete, terminally redundant viral genome (Supplementary Fig. S1).

Euryarchaeota comprise a large, diverse archaeal phylum that, in addition to MGII, includes thermophiles, halophiles and methanogens (*16, 17*). Viruses have been previously isolated from members of different euryarchaeal groups. The morphologies of euryarchaeal dsDNA viruses are diverse and include spindle–shaped, unclassified icosahedral, pleomorphic and head-tailed viruses. The latter group resembles the bacterial head-tailed phages (order *Caudovirales*), both in morphology and genome organization, and is currently classified into the families *Myoviridae*, *Podoviridae* and *Siphoviridae*, each of which contains bacterial and archaeal viruses (*18-20*). Head-tailed archaeal viruses so far have been found to infect only Euryarchaea, including mainly halophiles and methanogens (*21-23*). No viruses infecting MGII have been reported so far (*18*), and despite the increasing amount of viral metagenomic data released in recent years (*24, 25*), candidate viral contigs from the MGII group have not been reported either.

The protein sequence of the predicted MCP encoded in MAG 156409 showed significant, albeit moderate (33% identity), similarity solely to the MCPs of haloarchaeal siphoviruses (haloviruses) (Fig. 1A). Three additional proteins encoded in MAG 156409, namely, primase (Fig. 1C), portal protein and prohead protease (Supplementary Fig. S2 and S3), showed comparable levels of similarity to homologs from haloviruses, indicating a relatively distant evolutionary link. We used these protein sequences as queries in BLASTP searches against our entire Red Sea assembly dataset, in an attempt to expand the repertoire of virus-related sequences for subsequent, comprehensive comparative genomic analysis. This search resulted in additional 9 MAGs (Supplementary Table 1), all showing the same pattern of homology to haloviruses for all tested genes.

**Figure 1:**
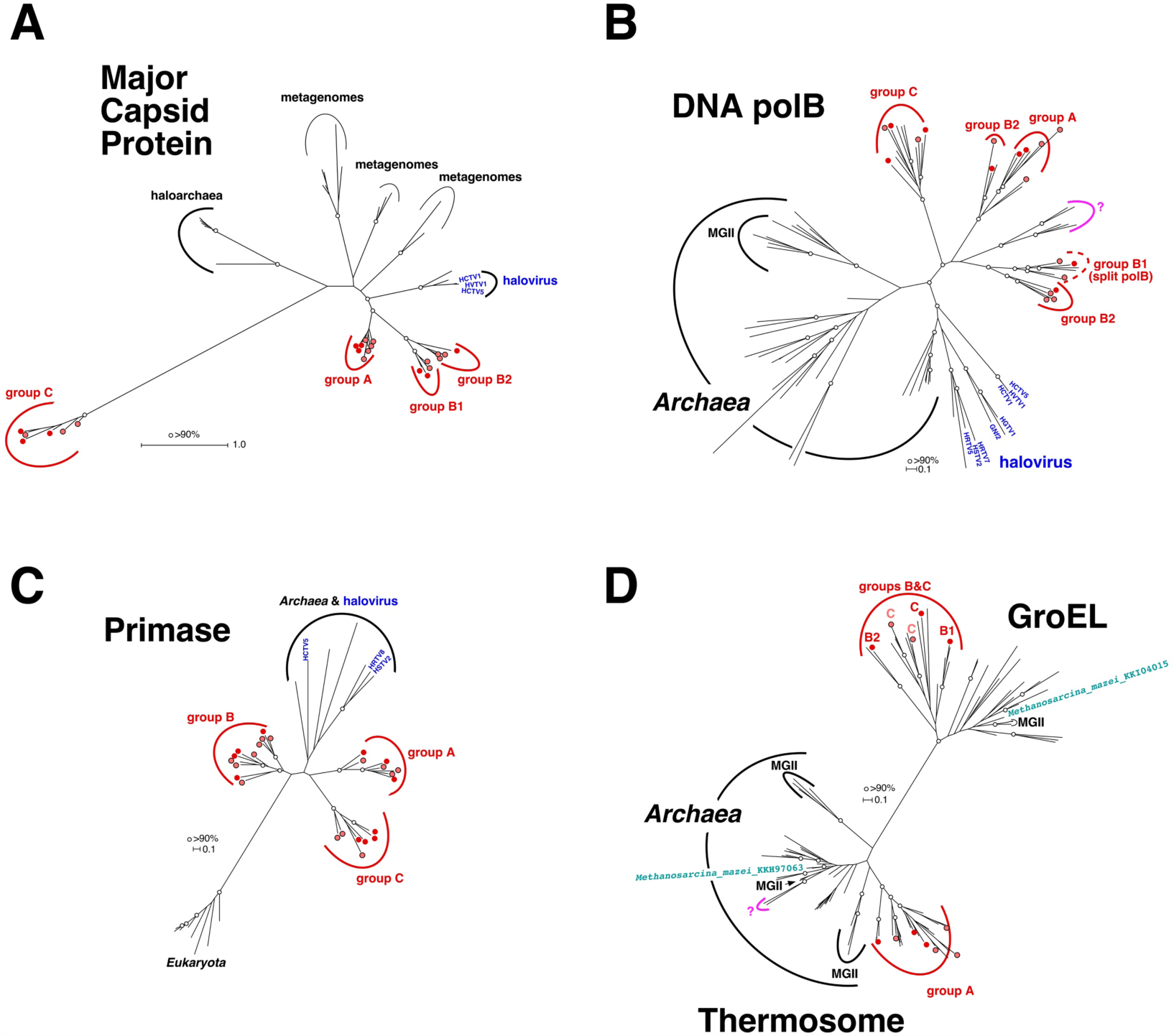
Unrooted Maximum likelihood phylogenetic trees of conserved Magrovirus genes. (**A**) Major Capsid Protein (MCP); (**B**) DNA Polymerase B (DNAP); (**C**) archaeo-eukaryotic Primase (AEP); (**D**) Chaperonins (thermosome subunit and GroEL). Metagenome-assembled complete or nearly complete genomes (MAGs) from the Red Sea and *Tara* Oceans metagenomes are marked with red and light red circles, respectively. Bootstrap support values greater than 90 are marked with white circles.

To validate and expand our observations on Red Sea metagenomes, we used the same query sequences in a BLAST search against both the original *Tara* Oceans assembly datasets (*26, 27*) and a reassembly of the raw data from this project (Sharon, in prep. see methods). This *Tara* Oceans search yielded 15 additional putative viral genomes related to the Red Sea MAGs (Supplementary Table 1). All together, we identified 26 putative viral MAGs that were assembled from two independent metagenomic projects (Red Sea and *Tara* Oceans).

In the phylogenetic tree of the euryarchaeal virus and provirus MCPs, and their environmental homologs (Fig. 1A), the MAG proteins split into 3 distinct groups, two of which (A, B) join in a clade affiliated with the halovirus MCPs, whereas the third group (C) forms a long branch with an uncertain affiliation. A similar phylogenetic pattern was observed for other hallmark caudoviral genes of the MAGs, namely prohead protease (Supplementary Fig. S3), portal protein (Supplementary Fig. S2), and large subunit of the terminase (Supplementary Fig. S4). Whereas groups A and B cluster together in all these phylogenetic trees, the position of group C changed from tree to tree, suggesting rapid evolution. These findings, along with the fact that 11 MAGs are terminally redundant linear genomes of about 100 kbp in size (Supplementary Table 1, Supplementary Fig S1), suggest that these MAGs represent a novel family of head-tailed archaeal viruses. Because, as we show below, these MAGs appear to be strongly associated with MGII, we provisionally denote them Magroviruses (<u>MA</u>rine <u>GRO</u>up II viruses).

Having tentatively delineated 3 distinct Magrovirus groups from the hallmark gene phylogenies, we compared their gene complements and genome organizations in detail. Although, as with many other archaeal viruses (*28*), most of the Magrovirus genes encode proteins without significant similarity to any sequences in current databases, the genomes include two readily identifiable, compact gene blocks, the structural-morphogenetic module and the replicative module (Figure 2A). The structural module consists of the genes encoding major capsid protein, portal protein, prohead protease, terminase, and closely resembles the corresponding genomic module of haloviruses (Figure 2B). The distinctive feature of Magroviruses is the presence of a suit of genes for proteins involved in the genome replication (Figure 2B). All 26 genomes encode DNAP, sliding clamp, clamp loader ATPase (replication factor C), archaeo-eukaryotic primase (AEP), replicative helicase (MCM protein), RadA-like ATPase, single-stranded DNA-binding protein (ssb), and several nucleases; all viruses except for Group C also encode one or two ATP-dependent DNA ligases (Figure 2B). Taken together, these proteins could comprise a nearly complete archaeal-type replisome (*29*), which is unprecedented among currently known viruses of archaea and bacteria. In particular, haloviruses that share the morphogenetic gene block with the Magroviruses encompass a smaller complement of replicative genes (Fig. 2 and Supplementary Fig. S5). The gene order within both the replicative and the structural modules of Magroviruses is highly conserved, with limited rearrangements, largely in group C (Figure 2B), possibly, because operonic organization of functionally linked genes is important for virus reproduction.

**Figure 2:**
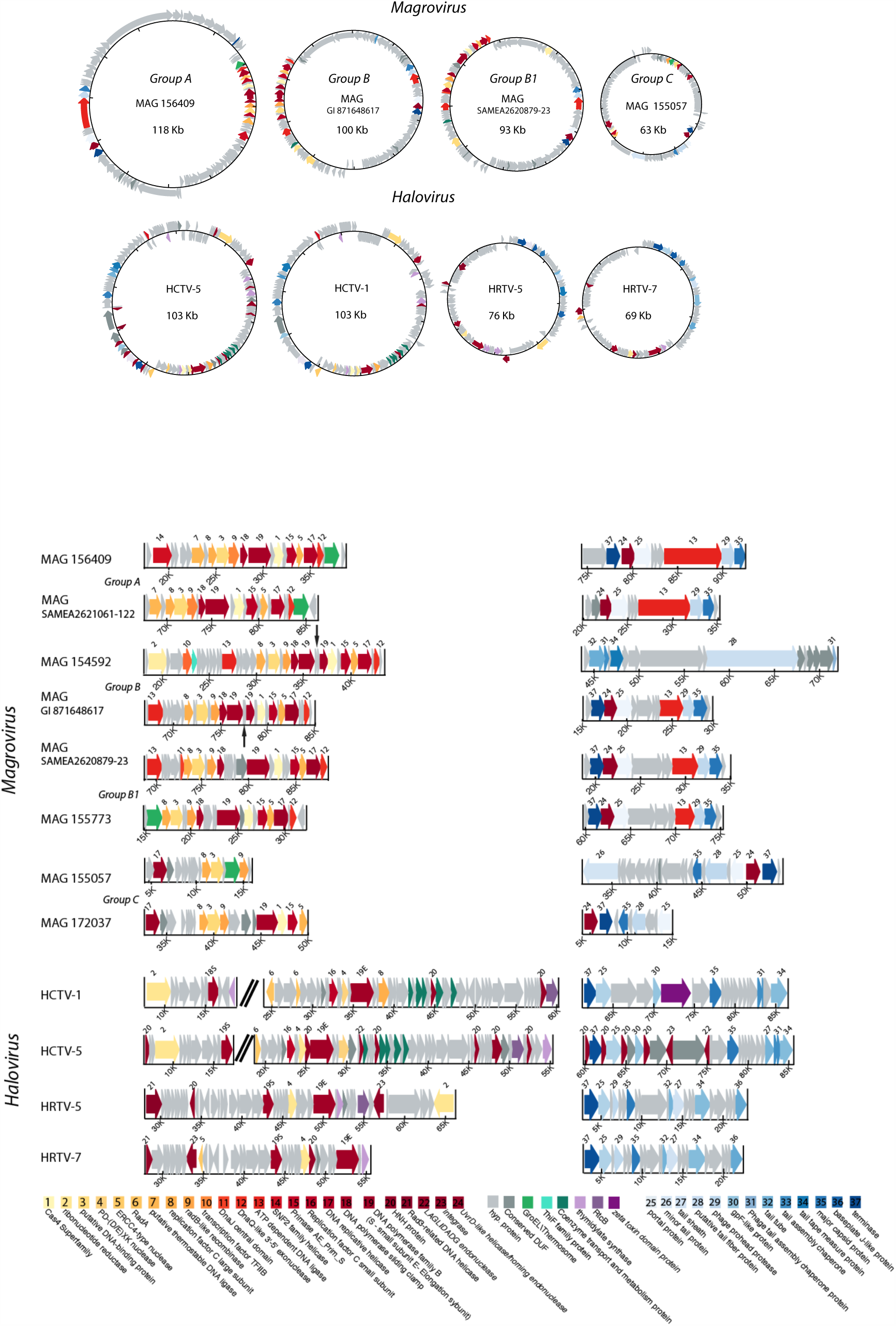
Genome organization in different groups of Magroviruses and haloviruses. For each group, the overall genome organization is shown (**A**) followed by detailed schemes (**B**) of the replicative gene block (left, yellow to red) and the structural gene block (right, different shades of blue). Homologous genes with predicted functions are shown using color code (see key in the bottom). Green arrows indicate thermosome genes and gray arrows indicate hypothetical proteins. Split DNAP genes from group B1 are marked with a narrow black arrow pointing to the regions between the split genes. See Supplementary Fig. S5 for greater details.

With the exception of the DNAP, AEP and ligases, the replicative proteins of Magroviruses show low sequence similarity to their homologs from cellular organisms that could be detected only through sensitive profile-profile searches. Nevertheless, most of these proteins show signs of archaeal origin as indicated by the provenance of the closest homologs (Supplementary File S1). Due to the low sequence conservation, informative phylogenetic analysis was feasible only for DNAP and AEP. In the phylogenetic tree of DNAPs, the Magrovirus polymerases cluster with the halovirus DNAPs (*23, 30*), and together, these viral polymerases are related to archaeal DNApolB III that is involved in lagging strand replication (Figure 1B). This gene is apparently subject to frequent horizontal transfer among archaea, so that its phylogeny does not follow the archaeal evolutionary tree. It appears likely that the transfers of the PolB III genes were at least partially mediated by viruses (*31*). Groups A and C that were delineated by analysis of morphogenetic genes retain monophyly in the DNAP tree (Figure 1B). In contrast, group B splits between two branches one of which is affiliated with group A (Figure 1B). This clear discrepancy between the MCP and DNAP phylogenies suggests genetic exchange among diverse Magroviruses. In addition to groups A, B, and C, a putative new group (labeled “?” in Fig. 1B & D) was observed in the DNAP tree. The MAGs encoding this group of DNAPs (tentatively, Group D) are not represented in the Red Sea metagenomes but are present among scaffolds originating from the *Tara* Oceans project. Group D genomes contain genes for DNAP (Fig. 1B), thermosome (Fig. 1C), resolvase, and DNA-dependent RNA poymerase A’, all with the highest similarity to archaeal homologs (Supplementary File S2). So far no closed genomes of this group were assembled, and a morphogenetic gene block was not identified. Therefore, although a connection to Magroviruses is apparent, the true nature of group D (virus, provirus, mobile element, megaplasmid, or uncultured archaea) remains to be determined once complete genomes become available.

Notably, 7 group B Magroviruses possess a split DNAP gene whereas in all other viral MAGs the DNAP gene is uninterrupted (Fig 2B and Supplementary Figuers S5 and S6). The split in the group B DNAP gene is located within the sequence encoding the catalytic domain, similarly to the DNAP genes of the cyanophage P-SSP7 (*32, 33*) and *Methanobacterium thermoautotrophicum*. In the latter case, it has been directly shown that the two portions of the split DNAP are expressed and interact to form the active polymerase (*34*). Interrupted DNAP genes are also found in other Euryarchaeota in which the inserts consist of post-transcriptionally excised inteins (*35-37*). The positions of the split in these archaeal polymerase and the group B Margovirus DNAP are similar but not identical, suggesting independent, convergent evolutionary events resulting in gene fragmentation. Regardless of the exact evolutionary scenario, the split DNAP is so far unique among archaeal viruses and supports the monophyly of Magrovirus Group B.

The phylogenetic tree of the AEP supports the monophyly of all 3 groups of Magroviruses as well as the common origin of primases in Magroviruses and haloviruses; in this case, however, Magrovirus groups A and C cluster with haloviruses, suggesting the possibility of gene exchange between these archaeal viruses (Fig. 1C).

Except for Group C, all Magroviruses encode a DnaQ-like exonuclease that can be implicated in proofreading during viral genome replication. Unlike most of the other Magrovirus genes, this nuclease shows significant sequence similarity only to bacterial homologs (Supplementary Fig. S6). Thus, somewhat unexpectedly, Magroviruses appear to have acquired genes not only from archaea but also from bacteria.

A notable feature of Magroviruses is the presence of two genes encoding distinct ATP-dependent DNA ligases genes in groups A and B, one located within the replicative module and the other one, unexpectedly, embedded in the structural module (Figure 2B). The provenances of these ligases are quite different as indicated by phylogenetic analysis: the ligase encoded within the structural module represents a distinct family that along with the Magrovirus proteins includes ligases from uncharacterized bacteria; the ligase in the replicative block of Group B is a typical archaeal variety, whereas the one in Group A replicative block belongs to the distinct family known as ‘thermostable ligases’ (Supplementary Fig. S5). Thus, apparently, ligases have been acquired by Magroviruses on 3 independent occasions.

The structural modules of all Magroviruses also contain another inserted gene that in different groups encodes distinct nucleases (Figure 2B). In groups A, B, and D, this is a homing endonuclease (LAGLI-DADG family) homologous to nucleases of mobile elements, such as Group I self-splicing introns and inteins, which are also present in many bacteriophages. In contrast, in Group C, the inserted gene encodes a homolog of the exonuclease subunit of the archaeal DNA polymerase D (*38*). An intriguing possibility is that the two nucleases play analogous roles in Magroviruses replication. In addition to the conserved replicative genes, several genes implicated in replication are found in individual groups of Magroviruses, *e.g.* ribonucleotide reductase in Group B and SNF2 family helicase in Group A.

Apart from the replicative and structural-morphogenetic proteins, 12 of the 26 Magroviruses encode either a bacterial-type chaperonin GroEL (Groups B, B1 and C) or a thermosome subunit, the archaeal homolog of GroEL in Groups A and D (Fig. 2B). Unlike the replicative genes, these Magrovirus proteins are highly similar to the archaeal and bacterial homologs. Phylogenetic analysis confirmed the sharp split between GroEL and thermosome subunits (Fig. 1D). The thermosome subunits of Magroviruses group with homologs from MGII, which is compatible with relatively recent acquisition of these genes by the viruses. A subset of MGII archaea encode GroEL instead of the thermosome subunit, conceivably owing to displacement of the ancestral archaeal chaperonin by a bacterial homolog. Although the Magrovirus GroEL do not group with those of MGII in the phylogenetic tree, this could be due to the accelerated evolution in the viruses; acquisition of this gene from MGII archaea remains a likely scenario. Comparison of the topology of the chaperonin tree with those of the MCP, DNAP and AEP trees suggests that the common ancestor of the Magroviruses acquired a GroEL gene that was replaced by the thermosome subunit in Group A. Chaperonins are not encoded by any known archaeal viruses but have been detected in several bacteriophages. Notably, these phages do not encode the co-chaperonin GroES, and GroES is not required for the phage chaperonin activity *(39)*. This is likely to be the case for the Magrovirus chaperonins as well. Given that, in Magroviruses, the chaperonin genes reside in the replicative gene cluster, it seems plausible that the chaperonins facilitate folding of the replicative proteins and replisome assembly.

Mapping reads from the Red Sea and from the *Tara* Oceans on the 26 Magrovirus genomes indicate that these viruses are globally widespread, with higher abundance in the Indian Ocean, the Red Sea and the South Pacific and Atlantic Oceans (Fig 3B). Furthermore, Magroviruses are highly abundant in the marine environment, third only to SAR11 phages and cyanophages (*40*). Surprisingly, the overall abundance of Magroviruses was found to be higher than that of SAR116 phages, previously reported to be the second most abundant phage group in the oceans (*41*) (Fig. 3A and Supplementary Fig. S9).

**Figure 3:**
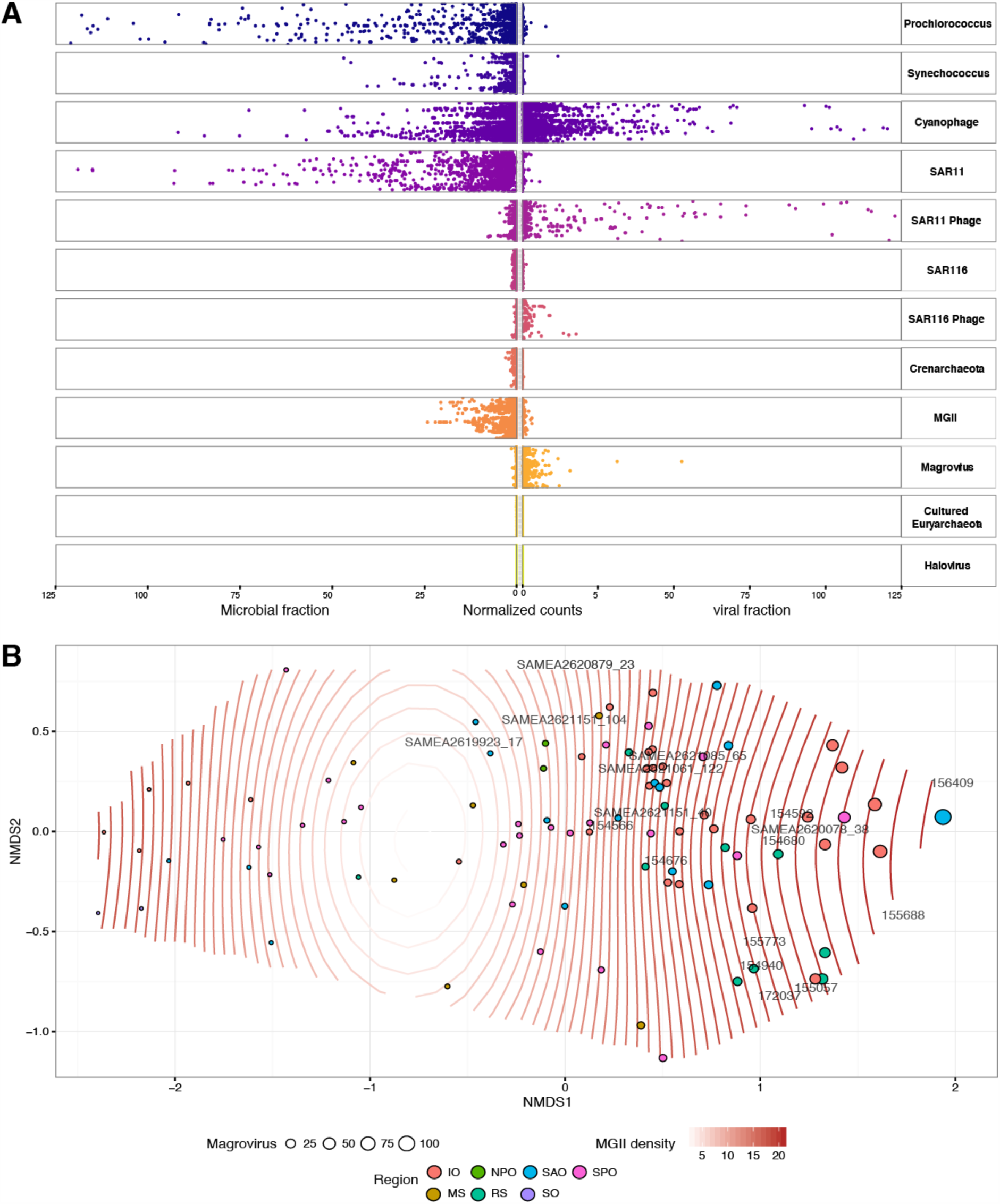
**A. The global spread of Magroviruses**. The abundance of the Magrovirus reads in Red-Sea samples and all *Tara* Oceans microbiome (*27*) and virome (*24*) is shown along with the abundances of the putative host MGII; other major archaeal groups; Cyanobacteria *Synechococcus* and *Prochlorococcus* and their phages; SAR11 bacteria and their phages; SAR116 and their phages. Horizontal axis: the normalized count (genome fragments per kilobase reference sequence per million library reads (GFPM), see methods.); vertical axis: the sampling stations. **B. NMDS Plot of sampling sites Magrovirus abundance**. circle diameter indicates Magrovirus abundance at a specific site. MGII abundance was fitted using *Ordisurf* function in Vegan package in R (*44*) and is represented by white to red contour lines. A linear response between the Magrovirus abundance ordination and the MGII abundance variable is represented by fitted contours that are equally spaced parallel lines perpendicular to the MGII abundance vector (R-sq.(adj) = 0.432 Deviance explained = 48%, p-value = 4.82e-10). Additional NMDS plots with smooth fitted surface for Cyanobacteria, Euryarachaeota and are shown in Supplementary Fig. 11. Region abbreviations: IO – Indian Ocean; NAO – North Atlantic Ocean; RS – Red Sea; SO – Southern Ocean; MS – Mediterranean Sea; NPO – North Pacific Ocean; SAO – South Atlantic Ocean; SPO – South Pacific Ocean.

So far, based on microscopic measurements (*30*), head-tailed viruses appeared to be the least abundant morphotype of archaeal viruses in the environment. However, our abundance estimates for Magroviruses suggest that these previously uncharacterized head-tailed viruses dominate the archaeal virome in surface marine environments. In contrast to the cyanophages, and to a lesser extent the SAR11 phages, the normalized counts of Magroviruses are almost negligible in the microbial fraction (larger than 0.2μm) of the samples but high in the virus-enriched fraction (<0.2 μm) (Fig. 3A). Thus, the principal source of the Magrovirus reads appear to be free virus particles rather than cell-associated viruses, proviruses or megaplasmids (Supplementary Fig. S8).

Despite the high abundance of MGII in the oceans and their apparent ecological importance, no viruses infecting these organisms have been identified so far. Here, using metagenomic approaches, we describe 3 distinct groups of viruses associated with MGII. Unequivocal demonstration of the virus-host relationship between magroviruses and MGII is currently unfeasible due to the lack of cultivable MGII representatives. Nevertheless, several lines of evidence strongly suggest that MGII archaea are indeed the hosts of Magroviruses. First, our abundance estimates show that MGII is the dominant archaeal group in the samples from which the Magrovirus genomes were assembled. Moreover, our Red-Sea assembly dataset consisted almost exclusively of MGII and Magrovirus contigs (Fig 3A). Second, most of the replicative genes of Magroviruses and especially the viral chaperonins show clear signs of archaeal provenance, and in some cases, a specific connection with homologs from MGII. Furthermore, Group B Magroviruses encode two tRNAs (tRNA^Leu^ and tRNA^Arg^; Supplementary Fig. S10) that are most similar to the respective tRNAs of MGII archaea.

The discovery of the Magroviruses is part of the growing trend in virology whereby viruses are recognized solely from metagenomics sequence analysis, a practice that has been recently formalized by the International Committee for Taxonomy of Viruses (*42*). Genome analysis of the Magroviruses is consistent with modular evolution of viruses whereby the structural and replicative modules have distinct origins. Magroviruses are unusual among viruses of bacteria and archaea in that they encode an elaborate, apparently (almost) self-sufficient replication apparatus. The closest analogy are the large and giant viruses of eukaryotes in the proposed order “Megavirales” that replicate in the cytoplasm of eukaryotic cells (*43*). Given the high abundance of both MGII archaea and Magroviruses, the latter are likely to be important ecological agents, similar to cyanophages and members of the “Megavirales” that infect abundant unicellular algae.

## ACKNOWLEDGMENTS

The authors would like to thank the captain and crew of the R/V *Sam Rothberg* of the Inter University Institute in Eilat, Israel, for their expert assistance at sea and Hagay Enav and Idan Bodaker for their help with sampling. The authors would also like to thank David Cohen of the Physics Department at the Technion for his help with the HPC ATLAS cluster. This work was partially funded by a European Commission ERC Advanced Grant (no. 321647), the People Programme (Marie Curie Actions) of the European Union’s Seventh Framework Programme FP7/2007-2013/ under REA Grant Agreement no. 317184, the Technion’s Lorry I. Lokey Interdisciplinary Center for Life Sciences and Engineering and the Russell Berrie Nanotechnology Institute, and the Louis and Lyra Richmond Memorial Chair in Life Sciences (O.B.). N.Y. and E.V.K. are supported by intramural funds of the US Department of Health and Human Services (to the National Library of Medicine).

